# Serum extracellular vesicle depletion processes affect release and infectivity of HIV-1 in culture

**DOI:** 10.1101/081687

**Authors:** Zhaohao Liao, Dillon C. Muth, Erez Eitan, Meghan Travers, Elin Lehrmann, Kenneth W. Witwer

## Abstract

Extracellular vesicles (EVs, including exosomes and microvesicles) are involved in intercellular communication in health and disease and affect processes including immune and antiviral responses. Ultracentrifuged serum is depleted of EVs and, when used in culture media, reduces growth and viability of some cell types. In this study, we examined the effects of serum EV depletion processes on HIV-1 replication in primary cells and cell lines, including two HIV-1 latency models. Increased HIV-1 production was observed in certain EV-depleted conditions, along with cell morphology changes and decreased cell viability. Add-back of ultracentrifuge pellets rescued baseline HIV-1 production. Primary cells appeared to be less sensitive to EV depletion. ACH-2 and U1 latency models produced more HIV-1 under EV-depleted conditions, while virus produced under processed serum conditions was more infectious. Finally, changes in cellular metabolism and gene expression were associated with EV-depleted culture. In conclusion, the EV environment of HIV-1 infected cells has a substantial effect on virus production and infectivity. EV-dependence of cell cultures should be examined carefully along with other experimental variables. However, EVs may not be the only particles depleted by ultracentrifugation or other processes. Effects of EVs may be accompanied by or confused with those of closely associated or physically similar particles.

## INTRODUCTION

Extracellular vesicles (EVs) are a diverse group of bilayer-membraned particles that include so-called “exosomes” (canonically defined as budding into multivesicular bodies (MVBs) and being released upon MVB fusion with the plasma membrane) and “microvesicles” (often described as blebbing directly from the plasma membrane)^1,2^. The mode diameter of EVs in circulation approximates that of retroviral particles^1^, and EVs and retroviruses share many common features^3^. Indeed, HIV has been called a “Trojan exosome,” eluding the host immune responses in part by masquerading as an EV^3^.

The relationship between EVs and HIV-1 infection is an area of active study, with some contrasting findings. While several other viruses can replicate via viral genomes packaged into host EVs^4,5^, HIV-1 does not appear to be capable of transmitting infection through this route^6^. However, EVs produced by HIV-1-infected cells contain fragments of viral RNA^7^ and viral proteins such as Nef^8^ and Gag^9^ (although another study did not find Nef to be associated with EVs^10^). HIV infection may alter the number and size of EVs as well as the host microRNA and proteins contained in EVs, which in turn may have implications for immune activation and HIV-1 pathogenesis^11–14^. In the setting of HIV-1 infection, EVs containing viral or host components may contribute to or exacerbate other conditions, such as HIV-1- or opiate-mediated neuron damage^15,16^. Whether specific EVs enhance or oppose HIV infection remains unclear and likely is context-dependent. EVs from HIV-infected cells can facilitate infection by several different mechanisms: by forming aggregates that include and deliver HIV-1 virions^17^; by activating CD4+ T lymphocytes, rendering them permissive for HIV-1 infection^18–20^; and by activating latent HIV-1 infection^21^. On the other hand, EVs from CD4+ T-cells can act as decoys to prevent infection of cells^14^, while EVs derived from human semen appear to inhibit HIV-1 replication and transmission^22,23^.

We previously showed that many cell types grow more slowly in media prepared with serum that had been ultracentrifuged to remove EVs^24^. Serum EV depletion has been observed to alter cell migration^25^, and macrophages become more proinflammatory when grown without serum EVs^26^. In general, we observed a slight but significant decline in replication and viability in EV-depleted conditions^24^. The magnitude of these effects was variable, and, notably, an astrocyte cell line (U87) did not appear to be affected. Adding isolated EVs back to the EV-depleted medium rescued cell growth, suggesting that the reduction in cell growth was in fact due to the depletion of EVs (although a role for EV-associated or otherwise co-purifying factors cannot be ruled out). We also found that the majority of the EVs that were internalized by cells, in a protein-dependent fashion, were targeted to lysosomes^24^. The identity of any specific growth-promoting factors contained in the EVs, such as RNA, proteins, or lipids, remains unknown, as does the extent to which these putative factors are involved in nutrition, signaling, and/or information exchange.

Prompted by these findings, we sought to determine whether serum EVs might affect HIV-1 replication in vitro. We used serum depleted of EVs by two methods and examined the effects of media prepared with these sera on the growth and behavior of cells that are susceptible to or infected with HIV-1. We also assessed the influence of EV depletion on virus production and infectivity. Finally, possible cellular explanations for these results were probed. We conclude that two separate serum EV depletion protocols have a profound effect on HIV-1 replication and infection, and that EVs and/or closely associated or co-depleted factors tend to exert a virus-suppressive effect.

## METHODS

### EV Depletion

For “UC-EVD” FBS, FBS diluted 1:4^27^ with Dulbecco’s PBS or base culture medium was centrifuged in a Beckman ultracentrifugation tube at 110,000 × g for 18 hours at 4°C (*AH-629 rotor, k factor=242).* Supernatant was gently removed from the top down by pipette, avoiding disturbing the bottom of the tube, and used for preparation of EV-depleted media, which was then filtered through a 0.22 *μ*m filter (Millex no. SLG5033SS). “TF-EVD” medium was prepared with Gibco™ Exosome-Depleted FBS (Thermo Fisher, USA, Catalog # A2720801), depleted by the manufacturer using a proprietary method. “EVR” refers to EV replete medium.

### Single particle tracking analysis

Extracellular particle concentration was measured using a NanoSight NS300 (Malvern, Worcestershire, UK) equipped with a 532 nm laser or a NanoSight NS500 with a 405 nm laser. At least five 20 second videos were recorded of each sample at a camera setting of 10, and files were analyzed at a detection threshold of five using NanoSight NTA software version 3.1.

### Western Blot

Samples were lysed with RIPA buffer (Thermo Fisher, #89900). Protein concentration was determined by bicinchonic acid assay. 100 *μ*g of protein from each sample was loaded onto a Criterion 10% Tris-HCl gel (BioRad, Hercules, CA, cat #3451018) and electrophoresed. The proteins were then transferred to a PVDF membrane (BioRad, cat# 1620174) and blocked with 5% powdered milk (Bio-Rad, cat #1706404) in Dulbecco’s PBS (Thermo Fisher, cat# 14190250) +0.1%Tween^®^20 (Sigma-Aldrich, St. Louis, MO, cat # P9416) for 1 h. The membrane was subsequently incubated with mouse-anti-human CD63 primary antibody (BD, cat #556019), at a 1:1000 dilution for 1 hour, and mouse-anti-human CD81 (Santa Cruz Biotechnology, cat# sc-166029), at a 1:1000 dilution for 1 hour. After washing the membrane, it was incubated with a goat-anti-mouse IgG-HRP secondary antibody (Santa Cruz Technology, cat #sc-2005) at a 1:10,000 dilution for 1 h. The membrane was then incubated with a 1:1 mixture of SuperSignal West Pico Stable Peroxide solution (Thermo Fisher, cat# 34080) and Luminol Enhancer solution (Thermo Fisher, cat# 34080) for 5 min. The membrane was visualized on Amersham Hyperfilm^TM^ ECL chemiluminescence film (GE Life Sciences, PA, cat # 28906839).

### Cell culture

**Cell lines.** H9 and PM1 (both T-lymphocytic); chronically HIV-1-infected T-lymphocytic (ACH-2) and promonocytic (U1) cells; and TZM-bl and HEK-293T cells were obtained from AIDS Reagent and ATCC. Cells were grown in complete RPMI medium (R10) prepared with EVR, UC-EVD, or TF-EVD FBS as well as 1 mM L-glutamine (Thermo Fisher, MA, USA; Cat # 25030081), 1000 U/mL, and 1 mg/mL Pen-Strep (Thermo Fisher, MA, USA; Cat # 15140148) and 10 mM HEPES (Thermo Fisher, MA, USA; Cat # 15630080).

**Primary cells.** Blood was obtained from healthy human donors under a university-approved protocol (JHU IRB # CR00011400). Within 15 minutes of draw, blood was diluted approximately 2:5 in PBS/5 mM ethylenediaminetetraacetic acid (EDTA)/2% EV-depleted FBS, loaded over room temperature Ficoll (GE Healthcare Biosciences, MA, USA; Cat # 17-1440-03) in SepMate^TM^ 50 mL tubes (StemCell, Vancouver, Canada; Cat # 85450), and centrifuged at 1200 × g for 10 minutes. The plasma/PBMC fraction was centrifuged at 300 × g for 8 minutes, and incubated in red blood cell lysis buffer at 37°C for 10 minutes. For monocyte-derived macrophages (MDM), PBMC were pelleted at 400 × g for 6 minutes, resuspended in cell culture media, and plated at 10^7^ cells per well in 6-well plates. Differentiation proceeded for seven days in the presence of macrophage colony stimulating factor (M-CSF) (R&D Systems, MN, USA; cat # 216-MC) as described previously^28^. CD4+ T-cells were isolated from PBMCs via EasySep^TM^ Human Naïve CD4+ T cell Isolation Kit (Vancouver, CA; Cat # 19155) (2). Purity was determined by flow cytometry on a BD Fortessa using a CD4+ antibody (Becton Dickinson, NJ, USA; Cat #562658). Interleukin-2 was added as a baseline stimulant at a concentration of 10 U/mL. Cells were activated 24 hours after isolation with phytohemagglutinin (PHA) at a concentration of 5 *μ*g/mL, diluted in culture medium (Sigma-Aldrich, MO, USA; Cat # 693839-1G).

### HIV-1 infection

HIV-1.Rf and .BAL stocks were generated from chronically infected H9 and PM1 strains, respectively, and stored at −80°C. For H9 and PM1 cultures, HIV.Rf and HIV.BAL were added at a concentration of 500 ng/mL (p24) and incubated at 37°C for 4-6 hours. Cells were then rinsed with PBS and spun twice at 400 × g for 5 minutes each. For primary macrophage cultures, HIV.BAL was added at a concentration of 200-250 ng/mL (p24) and incubated overnight at 37°C before rinsing twice with PBS.

### Cell Viability

Cell viability was assessed using the Muse™ Cell Analyzer and the Muse™ Count and Viability Kit according to manufacturer’s instructions (EMD Millipore, Billerica, MA, USA; Cat # MCH100102) or the WST-1 cell viability assay (Roche, IN, USA; Cat # 5015944001). The plates were mixed on an Orbital Shaker on setting 2 (Bellco, NJ, USA; Cat # 7744-20220) at room temperature for 30 min and absorbance was measured at 630 nm using an iMark™ Microplate Absorbance Reader (Bio-Rad, CA, US). MTT cell viability assay (Thermo Fisher MA, USA; cat # V-13154) was performed by incubating culture plates with MTT reagent for four hours at 37°C, adding SDS lysis buffer, and incubating for four hours to overnight at 37°C. Absorbance was measured at 570 nm with a plate reader as above.

### HIV-1 p24 Assay

200 *μ*l of cleared cell culture supernatant from all evaluated samples was set aside for p24 assays (Perkin Elmer, Holland, via Thermo Fisher Cat # 50-905-0509) at appropriate dilutions and following the manufacturer’s protocol. p24 levels were determined based on the manufacturer’s p24 standard. All results represent at least three separate experiments.

### Luciferase Assay

Luciferase expression in TZM-bl, which contain a luciferase gene under the control of a retroviral LTR, was monitored following overnight exposure of cells to H9/HIV.Rf- or PM1/HIV.BAL-containing-media from infected cells. The luciferase assay was done according to the manufacturer’s protocol using the Luciferase Assay System (Cat # E1500; Promega, WI, USA) and was read on a Fluoroskan Ascent FL luminometer (Thermo Fisher, MA, USA).

### Flow Cytometry

Cells were removed from plates with gentle pipetting, washed with 1 × PBS, and stained for 20 minutes at room temperature in the dark. All antibodies were from BD (CD4, ApC-Cy7: Cat # 341095; CD18, FITC: Cat # 555923; CD106, PE: Cat # 555647; CD54, PE-Cy7: Cat # 559771; CD195, APC: 556903). Samples were washed with 2 ml of 1x PBS once, spun at 400 × g for 5 minutes, and resuspended in 300 *μ*l of PBS to remove excess antibodies. Using a BD LSRFortessa, PBMCs were gated on lymphocytes by forward and side scatter profiles. Data were analyzed using FlowJo software v10.1 (FlowJo, OR, USA).

### EV Add-back

Pellets from ultracentrifuged FBS were re-suspended and added at 1/200^th^ and 1/50^th^ of the re-suspension volume to separate wells containing EVR, UC-EVD or TF-EVD-serum-containing media. These pellets were co-added with virus stock at the time of infection.

### RNA Isolation and Gene Expression Analysis

HEK293T cells (5 × 10^5^) were grown on a 10 cm dish for 24 hours. Medium was replaced with fresh media containing EVR or UC-EVD serum. Following 48 hours of growth, RNA was extracted using Trizol (Trizol Reagent, Invitrogen) according to manufacturer’s instructions. RNA integrity was assessed by Agilent Bioanalyzer RNA 6000 Chip (Agilent, Santa Clara, CA), and 500 ng total RNA labeled according to the manufacturer’s instructions (Illumina TotalPrep RNA kit). Biotinylated aRNA (750 ng) was hybridized to Illumina Human HT12v4 bead arrays overnight, rinsed and incubated with streptavidin-conjugated Cy3. Arrays were scanned at a resolution of 0.54 microns (Illumina iScan), and intensities were extracted from the scanned images using Illumina GenomeStudio software V2011.1. Data were normalized by Z-Score transformation and analyzed with DIANE 6.0, a spreadsheet-based microarray analysis program. Z-normalized data were analyzed with principal component analysis (PCA). Z-Scores for paired treatment groups were compared using the Z-Ratio statistic to determine gene expression changes within each comparison. Expression changes for individual genes were considered significant if they met four criteria: Z-Ratio above 1.5 or below #x2212;1.5; false detection rate (FDR) < 0.30; a P-value statistic for Z-Score replicability below 0.05; and mean background-corrected signal intensity greater than zero. Gene set analysis was performed using the open-source DAVID Functional Annotation.

**miRNA qPCR Array.** Total RNA was harvested from ACH-2, U1, and MDM grown under the different conditions using the miRvana total RNA isolation protocol as previously described^29^. A custom TaqMan low density array was ordered from Thermo Fisher, containing qPCR assays for 47 common miRNAs and the snRNA U6. Reverse transcription (100 ng starting material for each condition), pre-amplification, TLDA card processing were done using Thermo Fisher/Life Technologies reagents per manufacturer’s protocol and as previously described^30^. Data were extracted and processed as previously described^30^, except that normalization was performed to the geometric mean of ten miRNAs detected in all samples (miRs-24, 17, 30b, 106a, 142-3p, 92a, 146a, 342-3p, 21, and 16). Hierarchical clustering (Pearson, average linkage) and visualization was done with MultiExperiment Viewer (MeV)^31^.

### Seahorse Assay

Macrophages were plated in a Seahorse 96-well plate and were differentiated for seven days (see above) in EVR and EVD media. Cells were washed three times with media that did not contain sodium bicarbonate. Following one hour of incubation in a 37 degree incubator, mitochondrial activity was assessed using the Seahorse XF96 Analyzer (Seahorse Bioscience)^32^ according to the manufacturer’s instructions. Briefly, oxygen consumption rate (OCR) was measured following sequential addition of 2 *μ*M oligomycin, 1 *μ*M FCCP, and 5 *μ*M rotenone/antimycin A.

### Data Availability

Gene expression data and miRNA microarray data have been deposited with the Gene Expression Omnibus (GEO) under accessions GSE89067and GSE88838, respectively, part of the SuperSeries GSE89068 and BioProject PRJNA350212. Any additional data are available upon request.

### Ethical Approval and Informed Consent

Primary human cells were obtained from blood donors under a healthy blood donor protocol approved by the Johns Hopkins Institutional Review Board (IRB # CR00011400). Blood products were obtained in accordance with all relevant guidelines and regulations. All donors provided informed consent.

## RESULTS

### EV depletion and effects on cell line viability and proliferation

FBS was processed by dilution and overnight ultracentrifugation (“UC-EVD”) as previously described^27^ or by a proprietary commercial process (Thermo Fisher or “TF-EVD”). These sera and unmanipulated FBS (“replete,” EVR) were used to make cell culture media as described in Methods. Relative abundance of particles in replete or depleted conditions was assessed by NanoSight nanoparticle tracking analysis (Figure 1A), revealing decreased particle concentration of approximately 60-70%, with the greatest particle depletion in the TF-EVD condition. However, we would like to make two important points. First, the apparent efficacy of ultracentrifugation-based particle depletion varies in our hands by lot of serum, ultracentrifuge run, and nanoparticle tracking instrument/protocol, ranging from approximately 50-70%, while the TF-EVD process efficacy ranged from about 70-95% (data not shown). Second, as the terminology indicates, single particle tracking does not distinguish EVs from other particles. In contrast, Western blot for EV markers CD63 and CD81 reveals that, while these markers are barely detected in unprocessed FBS, they are highly enriched in ultracentrifuge pellets, demonstrating EV depletion (Figure 1B). Similar results were obtained for CD9, but the ER marker calnexin, Golgi marker GM130, and nucleus marker nucleoporin were not detected in pelleted fractions (not shown). We conclude that both ultracentrifugation and the TF-EVD process significantly but variably deplete particles including EVs from FBS, and that the TF-EVD process appears to be more effective than ultracentrifugation.

**Figure 1.**
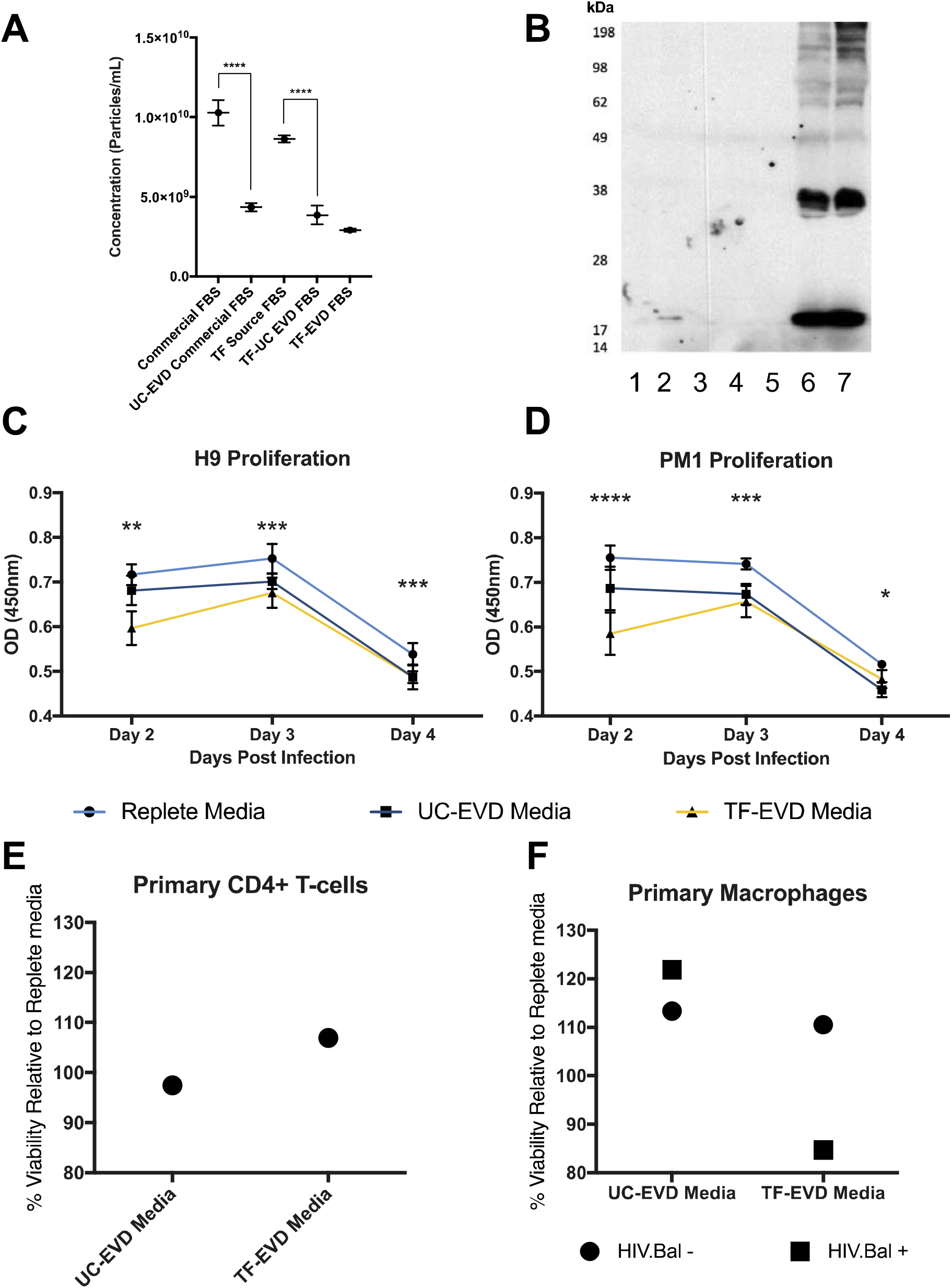
EVD conditions affect cell proliferation and viability. A) NTA of 25% FBS, 25% TF “source” FBS, UC-EVD of 25% commercial FBS, UC-EVD of 25% TF-Source FBS, and TF-EVD FBS and shows depletion of approximately 60% of particles by UC for both FBS types and the greatest depletion in TF-EVD FBS when compared with TF source FBS. Error bars represent standard deviation of 4 independent readings. **** p<0.005, one-way ANOVA with Tukey’s multiple comparison test. B) Western Blot analysis of 25% TF source FBS (1), 25% other commercial FBS (2), TF-EVD FBS (3), UC-EVD of 25% TF-Source FBS (4), UC-EVD 25% commercial FBS (5), and the UC pellet of TF Source FBS (6) and commercial FBS (7), and shows significant extraction of particles containing CD63 (MW ∼ 30-65 kDa) and CD81 (MW ∼22-25 kDa) C) For H9 cells, WST-1 assay optical density was greatest at all time points for the replete media conditions. D) PM1 cells: results similar to H9 cells. *=p<.005, difference between Replete and UC-EVD only; ** = p<0.005, differences between EVR and UC-EVD, UC-EVD and TF-EVD only; *** = p<.005, differences between EVR and UC-EVD and EVR and TF-EVD media; **** = p<.005, all types of media different from each other; Two way ANOVA with multiple comparisons. E) Viability of CD4+ T-cells at Day 7 post infection: cells grown in UC-EVD and TF-EVD media maintained similar viability to those in replete medium. F) MTT viability assay: no significant difference between infected, uninfected primary macrophages or between cultures grown in replete medium versus UC-EVD or TF-EVD depleted media.

Two T lymphocytic cell lines, H9 and PM1, were cultured in EVR, UC-EVD and TF-EVD media to assess the effect of EV-depleted serum on cell viability and growth. For all three conditions, WST-1 assay demonstrated small but significant (p<0.0001, 2-way ANOVA with multiple comparisons) differences in cell proliferation at Days 2, 3, 4 post initial culture conditions (Figure 1C, D) consistent with previous reports (23). Similar results were obtained using an LDH assay and by flow analysis with a Muse Cell Analyzer (data not shown). However, primary CD4+ T-cells and macrophages displayed no significant differences (Figure 1 E, F).

### Cells maintained in EV-depleted cell culture media exhibit increased HIV-1 production

T-lymphocytic H9 and PM1 cell lines cultured for seven days under the three conditions were infected with HIV-1.Rf and HIV-1.BAL respectively. Morphologic and behavior differences were observed in the TF-EVD condition when compared with the EV replete condition. Specifically, cells in TF-EVD medium tended to cluster and form syncytia (Figure 2A-C) more frequently than cells in EVR medium. Cells in UC-EVD medium were of intermediate phenotype that varied considerably by production lot of UC-EVD FBS, consistent with the variable and less efficient particle depletion by UC that we observed. Unexpectedly, HIV-1 release, as measured by p24 ELISA, was significantly increased in the TF-EVD condition in both H9 and PM1 cells (Figure 2D, E). Increased HIV-1 production was also observed from CD4+ T-cells and MDM infected with HIV-1.Rf and HIV-1.BAL strains, respectively (Figure 2F, 2G). Again, an increase was observed in many but not all UC-EVD conditions (data not shown).

**Figure 2.**
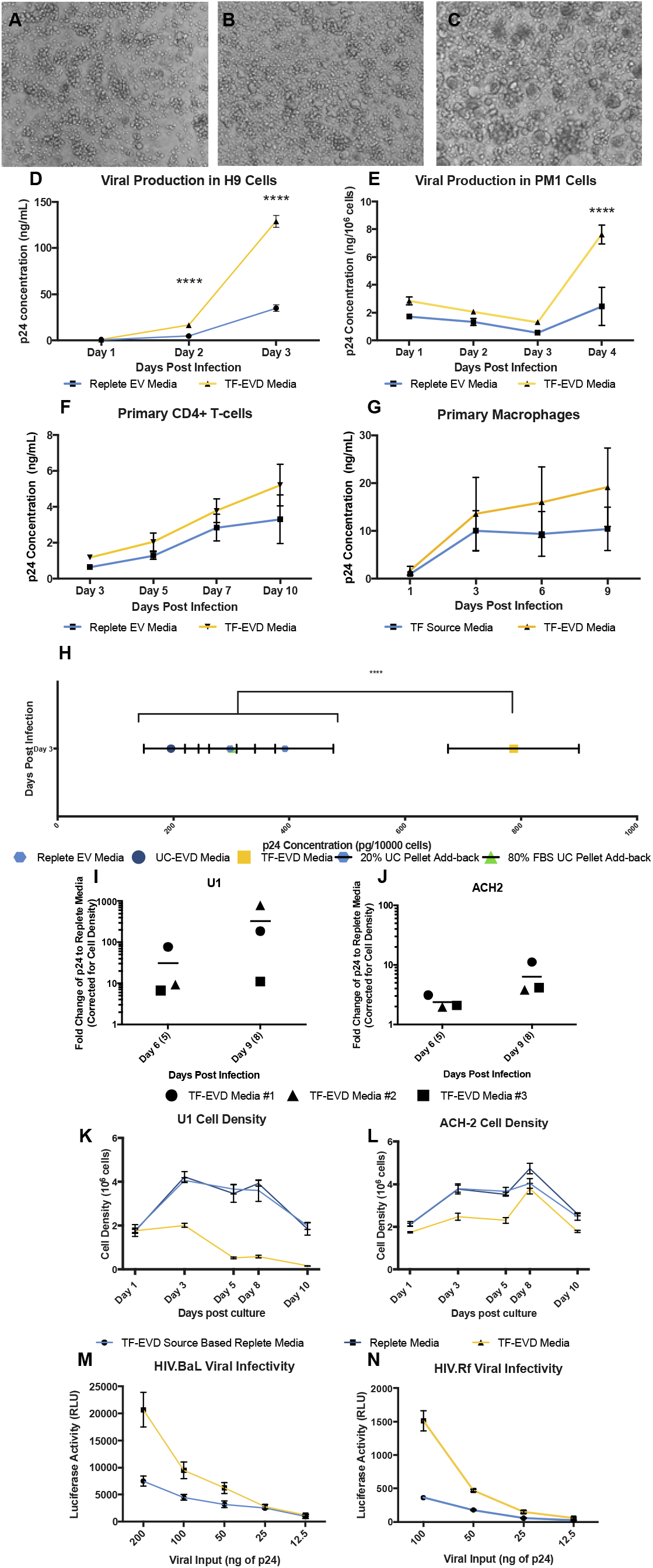
EVD serum affects cell morphology, virus production, and virus infectivity. Marked aggregation and syncytium formation was observed in extended culture of H9 T-cell line under EVD conditions (C) compared with EVR or UC-EVD conditions (A, B). D) HIV-1.Rf production in H9 cells is increased significantly in TF-EVD conditions by day 4. ****=p<.0001, 2 sample t-test with Holm-Sidak correction. E) Beginning at day 2, increased HIV.BAL production in TF-EVD conditions compared with EVR medium. ****: p<.0001; 2 sample t-test, Holm-Sidak multiple comparison correction. CD4+ T-cells infected with HIV-1.Rf (F) and monocyte-derived macrophages (MDM) infected with HIV-1.BAL (G) had a modest increase in virus production when grown in TF-EVD medium. H) Addition of previously removed EVs to EVD medium drops virus production back to near the EVR medium baseline. EVs isolated from stock FBS were added back to TF-EVD medium at 80% and 20% of their original concentrations. Supernatant was collected 1, 2, and 3 days post HIV-1.Rf infection. Measurements of virus release by p24 concentration at day 3 showed a pronounced difference between TF-EVD conditions and all others, ****=p<0.001, one-way ANOVA, Tukey’s multiple comparison test. I,J) Representative cell density data for U1 and ACH-2 cells from a single experiment. K,L) Representative experiments demonstrating that EV-depleted media produce virus with comparatively greater infectivity. HIV-1.BAL-infected PM1 Cells (M) or HIV-1.Rf-infected H9 cells (N) were cultured in EVR or TF-EVD medium conditions. Virus was collected at 7 days post infection and pelleted at 100,000xg for 2 hours. Pellet was diluted at 4 concentrations (shown on x-axis) in PBS and used to inoculate TZM/BL cells containing luciferase under the control of HIV LTR. After overnight incubation, an increase in luciferase activity was observed in the cells cultured with virus originating from TF-EVD medium compared with EVR medium at all concentrations.

### EV add-back restores HIV-1 production to baseline levels

To confirm that depleted material enriched in EVs contributes to the observed effects, resuspended EV-enriched UC pellets were added back to TF-EVD culture conditions. Irrespective of dose down to a 20% add-back, virus production was restored to levels significantly below the TF-EVD conditions (for example, HIV-1.Rf-infected H9 cells, Figure 2H). That virus production was not restored completely to baseline may be consistent with reports that UC-pelleted EVs may tend to aggregate and lose functionality^33–35^. Please note that, in the set of experiments depicted in Figure 2H, there was no significant difference between baseline and the UC-EVD condition, indicating the variability of the ultracentrifugation depletion technique.

### EV-depleted cell culture conditions reverse HIV-1 latency

ACH2 and U1 cells—models of HIV-1 latency in T-cells and monocytes, respectively—were cultured in replete medium and TF-EVD medium conditions. Significant differences in p24 production were observed in the TF-EVD condition as compared with the replete medium condition for both U1 and ACH2 cells in three experiments (Figure 2I, J). Cell density was correspondingly reduced after day one in both cell types (TF-EVD medium, Figure 2K, L). Of potential interest, the U1 cells were the most sensitive to depleted conditions of any cell type we studied. Nevertheless, the minority of live cells appeared to continue releasing relatively large quantities of p24. We do not think this p24 detection is simply the result of release from dead cells, as a similar phenomenon is not observed when cells are treated with toxins (data not shown).

### Heightened infectivity of viruses produced under depleted cell culture conditions

HIV-1 produced under depleted conditions was collected from H9 and PM1 cells and placed in p24-normalized amounts onto TZM-bl reporter cells, which encode a luciferase gene under the control of the HIV-1 LTR. Virus from TF-EVD conditions produced significant increases in luciferase expression compared with virus produced under EVR source serum conditions for all tested concentrations of virus (Figure 2M, N, S1).

### Assessing functional differences of cell lines raised in EVD conditions

To gather evidence for cellular changes that might possibly explain the observed responses to EV depletion, we performed several tests, including flow cytometry, respiration assays, miRNA qPCR array, and gene array.

**Flow cytometry: cell surface proteins.** The cell clumping observed in H9 cell cultures under EVD conditions prompted us to examine expression of several adhesion molecules and HIV-1 receptors by flow cytometry. Indeed, among other differences, VCAM-1 (CD106) and CCR5 expression were found to be increased on H9 and PM1 cells cultured in EV-depleted conditions (Figure 3A, B).

**Figure 3.**
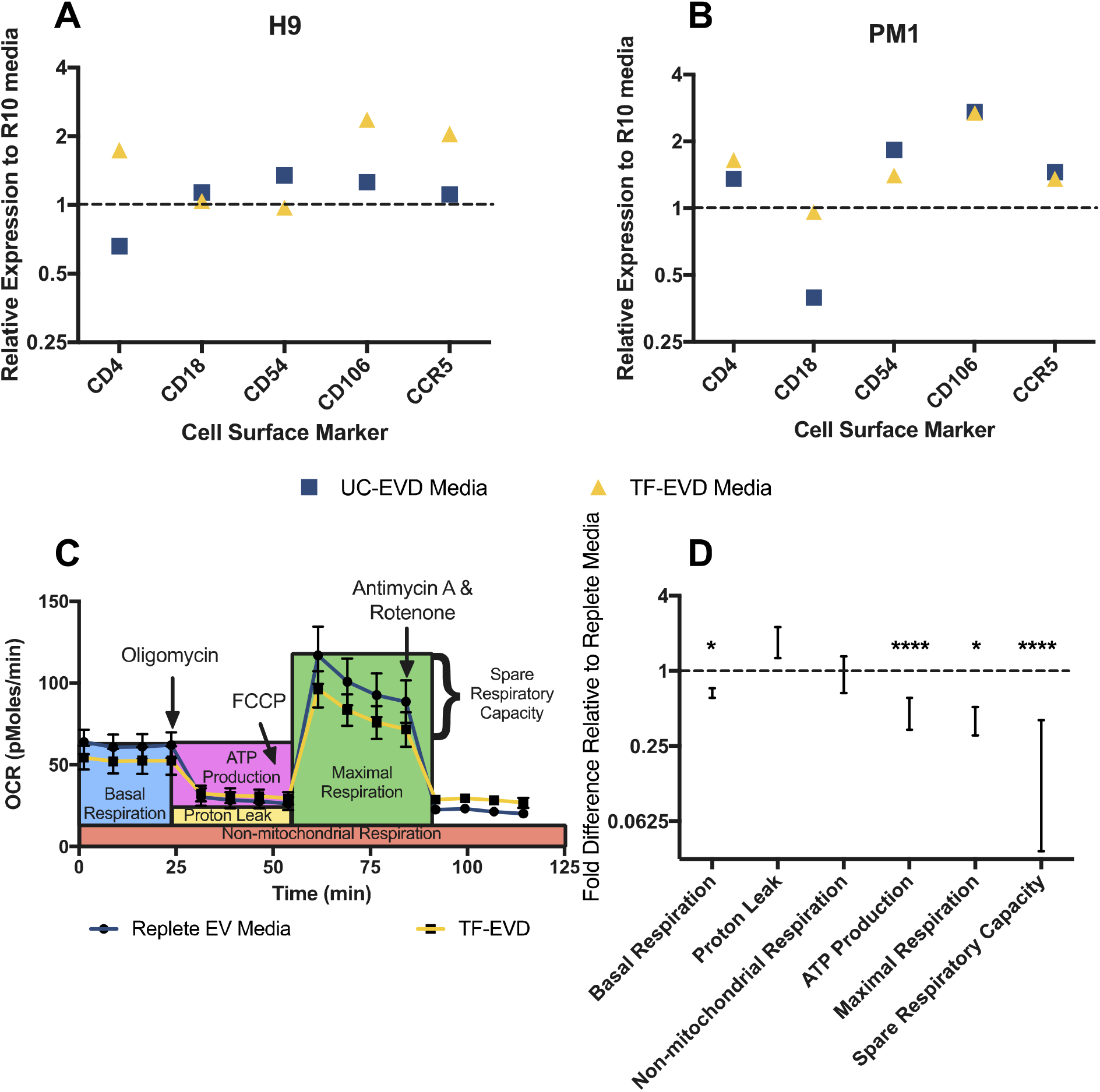
EVD conditions alter cell surface markers and respiration. A) H9 cells and B) PM1 cells displayed increased expression of selected surface/adhesion proteins by flow cytometry when cultured in UC-EVD or TF-EVD medium for 7 days; 1=expression on cells cultured in EVR. C,D) Oxygen compensation analysis of mitochondria of monocyte-derived macrophages grown in TF-EVD had significantly reduced basal and maximal respiration, as well as compromised ATP production and spare respiratory capacity. ****=p<0.001, *=p<.05, 2-way ANOVA with multiple comparison, Sidak test. Annotations in C) are adapted from the product materials (Agilent Technologies).

**Respiration assay.** We next used the Seahorse assay to test respiration under EV-depleted conditions (Figure 3C). Oxygen compensation analysis showed a significant, multiple-fold decrease in basal and maximal respiration, as well as compromised ATP production in mitochondria of MDM grown with EV-depleted media (Figure 3D). This reduction in mitochondrial respiration is in agreement with our previous and current observations of reduced cell growth.

**miRNA qPCR array.** Certain cellular microRNAs (miRNAs) have been reported to control retroviral replication^36,37^, while exogenous RNAs in cell culture medium are said to be taken up by cells^38^. We hypothesized that miRNA levels in cultured cells might be augmented by serum EV miRNAs, and that EV depletion might also indirectly affect miRNA expression. Either circumstance could result in reduced abundance of anti-HIV miRNAs, explaining an increase in HIV-1 production. However, results from miRNA profiling by a custom TaqMan low density array (TLDA) revealed no consistent changes in miRNAs across three cell types maintained in the different types of media. Unbiased hierarchical clustering showed cell type-specific miRNA expression patterns, but no indication of consistent differences associated with EVD conditions (Figure 4A). A total of five miRNAs appeared to be less abundant by 2-fold or more in one cell type under EV-depleted conditions (one in MDM, three in U1, one in ACH2), but no miRNA was 2-fold downregulated in more than one cell type, as one would expect if serum were an important and consistent source of miRNAs (Figure 4B). Importantly, previously reported anti-HIV miRNAs, including miRs-28-3p, −29a, −125b, −150, and −223, were not consistently diminished under EVD conditions. Similarly, there were no consistently upregulated miRNAs during EV depletion.

**Figure 4.**
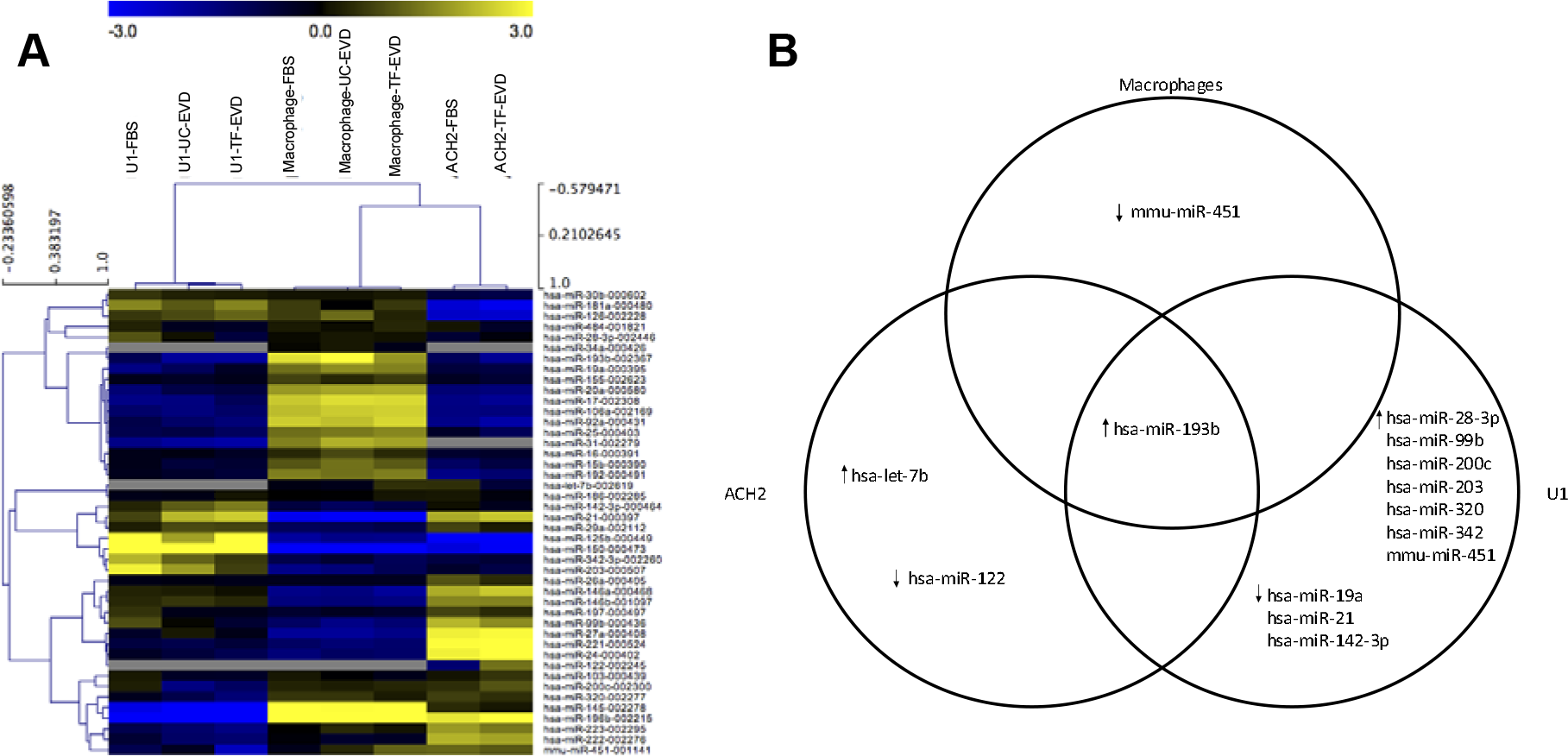
Little evidence for miRNA uptake from serum or serum-induced changes in miRNA expression. A) Unbiased hierarchical clustering demonstrates cell type-dependent miRNA expression, but no indication of differences associated with TF-EVD medium conditions. B) A total of five miRNAs appeared to be less abundant in TF-EVD conditions by two-fold or more in just one of the cell types, with no consistent findings between cell types. miR-193b was between 1.5- and two-fold upregulated in all three cell types.

**Gene expression.** Gene array analysis was performed with cells grown in EVR and EVD conditions. Gene ontology analysis of genes that were at least 2.5-fold more abundant in cells grown with EVD media revealed a significant increase in transcripts associated with lipid synthesis pathways, and especially sterol biosynthesis pathways (Table 1). It was previously reported that EVs contain a high proportion of cholesterol, sphingomyelin and ceramide; therefore, the increase in their biosynthesis may be a compensation response to loss of exogenous sources.

**Table 1.**
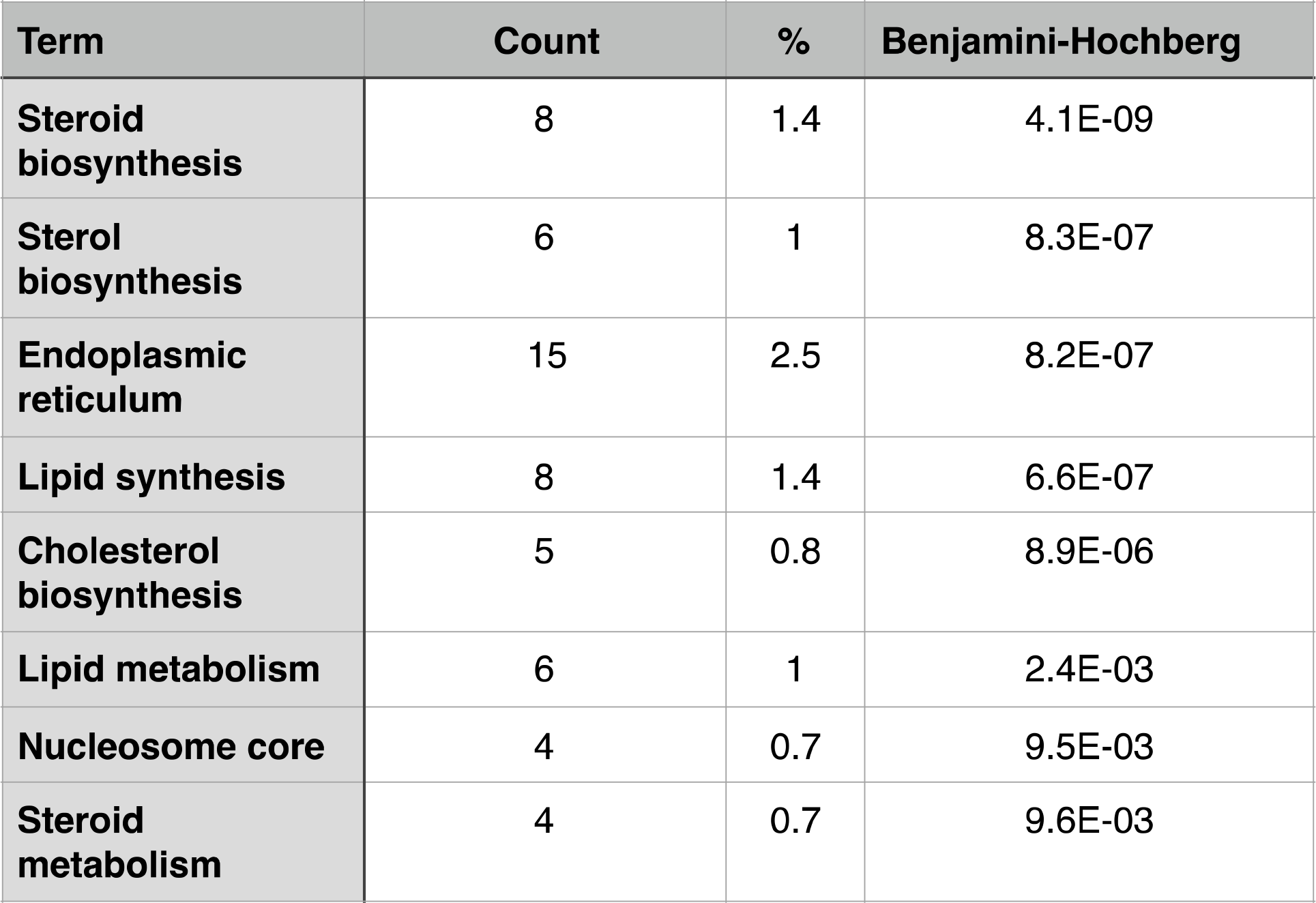
Gene Ontology Analysis reveals increased transcripts associated with lipid synthesis in EV-starved cells.

## DISCUSSION

We show here the significant impact of serum EV depletion protocols on virus production by HIV-1 susceptible cells. Additionally, HIV-1 latency models undergo a degree of viral activation under EVD conditions. Effects of EV depletion were less pronounced in primary cells, but we did observe a significant impact on HIV-1 production by primary macrophages. Altogether, our findings suggest that EVs can inhibit HIV-1 infectivity and release from leukocytes, and point to alterations in cellular lipids as a possible underlying mechanism.

Greater HIV production could be achieved through higher rates of infection (i.e., involving viral life cycle steps from cell binding through integration). The finding that certain cell adhesion molecules, including HIV-1 receptors and co-receptors, are upregulated on cells in EVD conditions might be consistent with more efficient infection. It could also explain the greater aggregation and syncytium formation we found in H9 cell culture. However, the observation that latency models are also affected by EVD conditions suggests that infection is not the sole explanation for our findings. Additionally, our EV add-back experiments—in which EV-enriched ultracentrifuge pellets were re-introduced into the medium simultaneously with virus exposure—suggest that any cellular responses to the absence of EVs would have been reversed rapidly, perhaps too rapidly to invoke receptor involvement. Alternatively, virion interactions with EVs or lipids that were added back could have inhibited infection (although this seems inconsistent with previous findings^39^).

Gene array data suggest another mechanism whereby virus production could be increased: in the relative absence of EVs, lipids involved in EV (and thus virion) biogenesis are upregulated. This may occur as the cell strives to achieve a homeostasis of membrane components, which must be sensed in some fashion. According to this hypothesis, normal interactions with diverse, non-self EVs or other lipid particles such as those found in serum preps could be viewed as a “security blanket” or a constant source of nutrition. In EV depletion, the cell might release its own EVs but be unable to make up for the presence of foreign entities. This is highly feasible, as culture media tend to contain much lower amounts of EVs than unprocessed serum. The observed surface upregulation of adhesion receptors is also consistent with this idea, bringing cells into closer contact with other membranes (cellular or EV). Notably, the sterol upregulation scenario would imply that the upregulation of HIV release is non-specific, or specific only to the extent that virions bud from sterol-rich membrane microdomains. To delve further into these possibilities, more information is needed on how cells sense the presence of EVs: through molecules of or on the EV surface, the cell surface, or both. One might also anticipate experiments to investigate the contribution of EVs from different cellular sources, which could explain the reviewed the differing findings in the literature reviewed previously.

Our latency model results are perhaps most exciting, suggesting that EV depletion could inform new strategies in HIV-1 eradication therapy. Depleting bulk EVs from plasma in vivo to increase HIV-1 production from latent cells in an eradication effort—in an approach analogous to leukapheresis—would be a difficult task. Even if such depletion were feasible, the same would likely be impossible in tissue. However, cells might be tricked into sensing a lack of EVs through pharmacologic means. Clearly, more knowledge about the system is needed to assess and develop this possibility.

As a final note, we would like to draw attention to an important caveat of this study. Although we concentrate here, in our language and experiments, on the effects of EVs, we hope that we have adequately conveyed that other interpretations are possible. The most accurate description of our results might be that putative EV depletion processes are associate with the observed effects. By extension, one might assume that EVs are involved, and this seems a reasonable assumption. After all, we can confirmed EV depletion by ultracentrifugation and by the commercial process, and the reversal of our findings during EV add-back is supportive. However, we cannot rule out that ultracentrifugation and other EV depletion protocols also deplete other components of serum. A recent publication from the Buzás lab found extensive low density lipoprotein contamination of EV preparations, and that LDL particles decorate EVs after co-incubation^40^. It is possible that LDL, other lipoprotein particles, and protein aggregates contribute to the phenomena we report here. We hope that others will join us in pursuing the many potentially informative studies that these findings might prompt.

## Acknowledgements and Funding

This work was supported in part by R01 DA040385 (to KWW), T32 OD011089, T32 GM008752, and seed funds from the Department of Molecular and Comparative Pathobiology (to KWW). The authors thank Thermo Fisher for providing Gibco™ FBS and Exosome-Depleted FBS for use in the experiments; Thermo Fisher had no role in study design, interpretation, or publication decisions. The authors are grateful to members of the Molecular and Comparative Pathobiology Retrovirus Laboratory for helpful discussions; to Mark Mattson and technical staff of the Laboratory of Neurosciences at the National Institute on Aging for support and discussion; and to Lisa Cimakasky for assistance with an early draft of the manuscript.

## Author Contributions Statement

KWW, ZL, and EE designed, planned, and supervised the study. ZL, DCM, EE, and MT performed experiments. EL performed gene array analysis and assisted with data submission. KWW and DCM wrote most of the manuscript. DCM assembled most figures. All authors reviewed and approved the manuscript.

